# Influenza Virus Infection of an Immunocompetent Organotypic Model of the Human Respiratory Mucosa

**DOI:** 10.64898/2026.03.24.713934

**Authors:** Annia Pérez-Riverón, Emma Deiss, Ambre Alléon, Pualani Ateni, Jianhui Li, Florent Foisset, Christine Lehalle, Jean-Daniel Fauny, Nelly Frossard, John de Vos, Redmond Smyth, Christian Debry, Léa Fath, Christopher G. Mueller, Benjamin Voisin, Vincent Flacher

## Abstract

Respiratory infectious diseases are among the leading causes of global morbidity and mortality and remain a major public health concern. Progress in understanding early host-pathogen interactions has been hampered by the limited physiological relevance of existing experimental systems. Different models mimicking the human respiratory epithelium have been developed to study viral infections *in vitro*, such as tridimensional (3D) tissue models and organoids. However, many lack key features of human tissue architecture, particularly the lamina propria or immune cells. To address these limitations, we established an immunocompetent 3D model of the human respiratory mucosa by combining nasal epithelial cells isolated from nasal brushings, fibroblasts from mid-turbinate nasal biopsies, and macrophages derived from blood monocytes. These cells were sequentially seeded into collagen-chitosan scaffolds, resulting in a reconstructed respiratory mucosa closely resembling the in vivo nasal tissues. To further confirm the physiological relevance of the model, we infected it with influenza A virus. The mucosa model supported viral replication in the epithelium and consequently showed increased secretion of inflammatory cytokines and upregulation of type I interferon related genes, enabling the monitoring of early antiviral innate immune responses in a physiologically relevant context.

## 1. Introduction

Respiratory infectious diseases represent a significant global healthcare burden and are one of the leading causes of death worldwide, with approximately 2.6 million deaths caused by lung infections annually^1^. They are also among the most common illnesses in children, with pneumonia representing the leading cause of death among those under 5 years old^2^. Moreover, highly pathogenic acute respiratory viral infections have caused six global epidemics and pandemics with high morbidity and mortality rates in the past 20 years^3^. Despite advances in epidemiological surveillance, diagnosis and preventive vaccination, respiratory infectious diseases continue to be a major global concern^4^, underscoring the need for innovative approaches to advance basic and translational research.

While animal models have greatly contributed to our understanding of respiratory disease pathogenesis, their applicability is limited by a low predictive value to human outcomes^5^. Together with the increased application of the 3R principle aiming to reduce, refine, and replace the use of animals in research, this has led to the development of novel research models that can faithfully recapitulate the essential physiological features of the human airways. Despite recent alternatives such as airway organoids or precision-cut lung slices, air-liquid interface (ALI) cultures of human bronchial epithelial cells have become a gold-standard to study the mechanisms of infection and pathogenesis of respiratory viruses^6,7^.

ALI cultures replicate the fully differentiated respiratory epithelium^8,9^, comprising different cell types such as basal, ciliated, goblet and club cells. This organotypic model preserves key tissue features like the production of mucus and cilia beating which are critical for studying host-pathogen interactions. ALI cultures are established by exposing the apical surface of the epithelium to the air while keeping the basolateral surface submerged in liquid medium. This configuration preserves tissue polarity, allowing physiologically relevant simulation of in vivo processes such as apical viral exposure and entry, as well as drug delivery^10–12^.

However, even though traditional ALI models can closely mimic the human airway epithelium, they are limited in reproducing the complexity of the respiratory tissue architecture. Consequently, there are growing efforts in the optimization of these models to include extracellular matrix (ECM) components^13,14^, stromal cells^6,14^ or immune cells^15,16^, all of which can strongly influence host-pathogen interactions. In particular, macrophages are among the most abundant immune cells in the airways^17^ and play key roles in immunosurveillance and tissue repair^18^.

Here, we report the development of an immunocompetent 3D tissue model of the human respiratory mucosa, comprising a fully differentiated nasal respiratory epithelium grown on a lamina propria equivalent that contains nasal fibroblasts and macrophages. We used this in vitro multicellular system as a platform to monitor early antiviral innate immune responses against influenza A virus (IAV).

## 2. Materials and Methods

### 2.1 Expansion of fibroblasts from mid-turbinate nasal biopsies

Tissue samples were collected from the mid-turbinate. Soft tissue was carefully separated from cartilage, minced, and added to 10mL of Dulbecco’s Modified Eagle Medium (DMEM) (Corning) containing 3mg/mL collagenase D (Roche), 1mg/mL DNase I (Roche), and 400μg/mL neutral protease grade II (Roche). The enzyme mix was incubated at 37°C under agitation (200 rpm) for 45min. The resulting cell suspension was passed through a 100μm cell strainer and centrifuged at 300LJ×LJg for 5min. Cells were resuspended in 25mL of PBS (Corning) containing 2% fetal bovine serum (FBS) (Corning) and 2.5mM EDTA (Sigma Aldrich), then passed through a 70μm cell strainer. After a second centrifugation at 300LJ×LJg for 5min, cells were counted and seeded at a density of 6×10^4^ cells/cm^2^ in DMEM medium supplemented with 10% FBS, 100U/mL penicillin/streptomycin (Corning), and 50μg/mL gentamicin (PAN Biotech). Cells were cultured at 37°C in a humidified atmosphere with 5% CO_2_ and passaged upon reaching 80% confluency. Medium was exchanged every 2-3 days.

### 2.2 Expansion of epithelial cells from nasal brushings

Samples were obtained by gentle brushing under general anesthesia during endoscopic surgery in patients undergoing procedures for benign tumors, following informed consent. Cell suspensions were collected in 5mL of PneumaCult-Ex Plus medium (Stemcell) and transported on ice. Suspensions were passed through a 70μm cell strainer and centrifuged at 300LJ×LJg for 5min. Pellets were resuspended in 2mL of EasyLyse™ Erythrocyte-Lysing Reagent (DAKO), previously diluted 1:20 in PBS and incubated during 2min at room temperature. To stop the lysis, 10-15mL of PBS were added, and samples were centrifuged again at 300LJ×LJg for 5min. Cells were seeded at a density of 25000 cells/cm^2^ in plates pre-coated with 25μg/mL of rat tail collagen (Sigma-Aldrich) with PneumaCult-Ex Plus culture medium. Cells were cultured at 37°C in a humidified atmosphere with 5% CO_2_ and passed when they reached 80% confluency, up to 3 passages. TrypLE Express Enzyme (Gibco) was used during 5min for detachment in each passage. Medium was exchanged every 2-3 days. The cells were phenotyped by immunofluorescence microscopy at different stages of the culture.

### 2.3 Purification of monocytes from human blood

Monocytes were isolated from human buffy coats obtained from the French National Blood Service (Établissement Français du Sang, Strasbourg, France). Peripheral blood mononuclear cells (PBMCs) were first isolated by density gradient centrifugation using Ficoll-Paque™ (Sigma-Aldrich). To this end, the buffy coats were diluted in PBS (50mL blood + 190mL PBS) and layered over Ficoll (15mL Ficoll + 35mL diluted blood), followed by centrifugation at 1200 × g for 20min at room temperature without brakes or acceleration. The PBMC layer was then collected, washed twice with PBS and centrifuged at 300 × g for 5min between washes.

For monocyte enrichment, PBMCs were subjected to a second density gradient centrifugation using Percoll (Sigma-Aldrich). The working solution was prepared by mixing 23.4mL Percoll with 2.6mL of 10x PBS and 26mL of PBS to achieve isotonic conditions. PBMCs were resuspended in PBS supplemented with 2% FBS and 2.5mM EDTA at a maximum concentration of 10^8^ cells/mL, and 3mL of this suspension was layered over 6mL of Percoll in 15mL tubes. Samples were then centrifuged at 1200 × g for 20min at room temperature without brakes or acceleration. The monocyte-enriched phase was collected, washed once with PBS supplemented with 2% FBS and 2.5mM EDTA and resuspended in the same solution. Monocyte enrichment rates were determined by flow cytometry from the proportion of CD45+ CD3− CD14+ cells. 70-80% pure monocytes were cryopreserved for downstream applications.

### 2.4 Differentiation of macrophages from blood monocytes

Purified human monocytes were seeded at a density of 1.6×10^5^ cells/cm^2^ in complete RPMI-1640 medium (Corning) supplemented with 10% FBS, 100U/mL penicillin/streptomycin, 50μg/mL gentamicin, and 50ng/mL macrophage colony-stimulating factor (M-CSF; Immunotools GmbH). Cells were cultured at 37°C in a humidified atmosphere containing 5% CO_2_. The medium was refreshed on days 3 and 5. On day 7, cells were detached by incubation with PBS supplemented with 2% FBS and 2.5mM EDTA at 37°C for 40min.

### 2.5 Flow cytometry

1-2×10^5^ cells were stained with directly fluorochrome-conjugated primary antibodies diluted in PBS supplemented with 2% FBS and 2.5mM EDTA for 30min at 4°C. Following incubation, cells were centrifuged at 300LJ×LJg for 5min and washed once with the same buffer. 0.5µg/mL DAPI (Sigma-Aldrich) diluted in PBS with 2% FBS and 2.5mM EDTA was added as a viability marker prior to acquisition. Flow cytometry was performed on a Gallios cytometer (Beckman-Coulter) and analyzed with FlowJo software (Becton-Dickinson). Gates were established based on the fluorescence minus one (FMO) negative controls.

The antibody panel used to phenotype purified monocytes included: mouse anti-human CD45-PE-Cy7 (Biolegend), mouse anti-human CD3-APC-Cy7 (BD Bioscience) and mouse anti-human CD14-PE (Immunotools). The panel used to characterize macrophages derived from blood monocytes included: mouse anti-human HLA-DR-AF700 (R&D Systems), mouse anti-human CD14-PE (ImmunoTools), mouse anti-human CD64-APC (BD Pharmingen), and mouse anti-human CD163-FITC (R&D Systems).

The panel used to characterize cells derived from nasal biopsies included: mouse anti-human CD45-PE-Cy7 (Biolegend), recombinant anti-human EpCAM-AF488 (R&D), mouse anti human-CD31-APC-Cy7 (Biolegend) and mouse anti-human CD90-PE (Biolegend).

### 2.6 Preparation of collagen-chitosan scaffolds

Type I bovine collagen (Symatese) and chitosan (CD Bioparticles) were dissolved in pre-warmed (37°C) 0.1% acetic acid at final concentrations of 10mg/mL and 2.8mg/mL, respectively. The collagen solution was incubated at 37°C under gentle rotation for ≥1h. The chitosan mixture was similarly agitated for ≥1h at room temperature, adding 2 drops of pure acetic acid every 10min until fully dissolved. The two solutions were combined by adding the chitosan to the collagen and incubating at 37°C with gentle rotation for ≥1h, followed by a brief centrifugation and a final 10min incubation.

Pre-warmed 12-well plates (37°C) containing Whatman paper rings (outer diameter 2.2cm; inner diameter 1.6cm) were filled with 500µL of the matrix per well using pre-heated cut tips. Plates containing the collagen-chitosan mixture were lyophilized (Aerial, Illkirch-Graffenstaden) according to previously published specifications^17^. 48h before utilization, plates were rehydrated and sterilized with 70% ethanol during 24h at 4°C. Ethanol was replaced by DMEM medium supplemented with 10% FBS, 300U/mL penicillin/streptomycin solution and 150μg/mL gentamicin. Plates were incubated 24h at 37°C and medium was replaced with DMEM with 10% FBS, 100U/mL of penicillin/streptomycin and 50μg/mL of gentamicin. Scaffolds were stored at 4°C until further use.

### 2.7 Generation of the nasal mucosa model

A multilayered tissue model was generated by sequential seeding of fibroblasts, macrophages, and epithelial cells onto collagen-chitosan scaffolds over a total culture period of 14 weeks. At each step, the cells seeded onto the scaffolds were incubated for 2h at 37°C in a humidified atmosphere with 5% CO_2_ before adding medium. Culture media was refreshed every 2-3 days throughout the entire protocol. Initially, 10^6^ nasal fibroblasts were seeded onto side 1 of the scaffolds in 100μL of DMEM/F12 medium (Gibco) supplemented with 10% FBS, 100U/mL penicillin/streptomycin, 50μg/mL gentamicin, 1x MEM non-essential amino acids (Gibco), and 5μg/mL of freshly prepared vitamin C (Sigma-Aldrich). After incubation, 4mL of the same medium was added per well. After 1 week, 5×10^6^ fibroblasts were seeded onto side 2 and cultured for 4 additional weeks in 4mL of medium. At week 5, 2.5×10^5^ macrophages were added to side 2 of the scaffolds in 100μL of medium, followed by the addition of 4mL per well. Control models did not receive macrophages. At week 6, 10^6^ epithelial cells expanded from nasal brushings were seeded onto side 1. After 2h, 5mL of PneumaCult-Ex Plus medium was added per well. Following 1 week of submerged culture, scaffolds were transferred to ALI conditions and cultured for 6 additional weeks in 3mL of PneumaCult-ALI medium (Stemcell) supplemented with 480ng/mL hydrocortisone (Stemcell) and 4μg/mL heparin (Stemcell) to promote epithelial differentiation. For each model, fibroblasts from at least two independent donors and epithelial cells from three independent donors were used.

### 2.8 Histology staining

Nasal mucosa models were embedded in paraffin and sectioned at 8μm thickness. Sections were then deparaffinized and rehydrated as follows: 10min in toluene, 2min in absolute ethanol, 1min in 95% ethanol, and 5min in Milli-Q water. Mucosa models were incubated overnight at room temperature in Bouin’s solution and, following fixation, washed under running tap water until the yellow coloration was completely removed. Sections were subsequently incubated for 5min in Weigert’s Iron Hematoxylin solution (Sigma-Aldrich), followed by a 5-min wash under running water and an additional rinse in Milli-Q water. Sections were then stained for 5min with Biebrich Scarlet-Acid Fuchsin solution (Sigma-Aldrich) and rinsed again in Milli-Q water. A phosphotungstic/phosphomolybdic acid solution was prepared at a 2:1:1 ratio of Milli-Q water, phosphotungstic acid, and phosphomolybdic acid and tissue sections were incubated in this solution for 5min. Slides were then transferred to Aniline Blue solution (Sigma-Aldrich) for 5min, followed by a 2-min incubation in acetic acid. After rinsing in Milli-Q water, sections were dehydrated in absolute ethanol during 1min, cleared for 10min in toluene, and mounted using Eukitt hydrophobic medium (Sigma-Aldrich).

### 2.9 Analysis of cilia beating frequency

Nasal mucosa models were placed in 27mm glass base dishes (Thermofisher) containing 2mL PneumaCult-ALI culture media with the apical side oriented to the glass. Differential interference contrast acquisitions were performed on a Zeiss Axio Observer Z1 microscope, equipped with environmental control and piloted by MetaMorph software (Molecular Devices). Acquisitions were conducted in stream mode. The exposure time depended on the size of the chip used, defining the acquisition frequency (ranging from 30 to 49 Hz). Images were processed using FIJI software^18^ and analyzed using the Fast Fourier Transform function from the NumPy library in Python. The first non-zero frequencies were retained for each pixel to extract cilia beating frequencies. Ilastik software^19^ was used for pixel classification to segment the regions described above, retaining only pixels associated with cilia. The mean frequencies were then measured for each image within these segmented regions.

### 2.10 Virus amplification

The influenza A(H3N2) strain A/Nepal/921/2006 was obtained from the International Reagent Resource (IRR; catalog FR-367). Madin-Darby canine kidney (MDCK) cells (FR-926 MDCK-ATL, IRR/CDC) were seeded at 3-6×10^4^ cells/cm^2^ and cultured in DMEM supplemented with 10% FBS and 100U/mL penicillin/streptomycin, and infected the following day when cultures reached approximately 80-90% confluency. Prior to infection, the monolayer was rinsed once with PBS with Ca^2^□/Mg^2^□ (Gibco) supplemented with 0.2% bovine serum albumin (BSA, BMD Millipore). The virus inoculum was gently added to the culture dishes at a multiplicity of infection (MOI) of 0.001-0.01 and incubated for 1h at 37°C, 5% CO_2_, with gentle rocking every 10min. After adsorption, the inoculum was removed, and cells were washed once with PBS with Ca^2^□/Mg^2^□. Cultures were overlaid with Opti-MEM maintenance medium (Thermofisher) containing TPCK-treated trypsin (Sigma-Aldrich) at 0.5-1µg/mL and incubated at 37°C, 5% CO_2_ for 36-72h. Supernatants were harvested when 50-80% of cytopathic effects (CPE) were observed or at 48-72h post-infection. Harvested supernatants were clarified at 500 × g for 15min at 4°C and filtered through 0.22µm polyethersulfone membranes. Titration of virus stocks was performed by plaque assay on MDCK cells as described below. The resulting virus titer was 1.94 × 10□ PFU/mL

### 2.11 Infection of reconstructed 3D nasal mucosa

Three days prior to infection, the medium in the scaffolds was replaced with PneumaCult-ALI medium without hydrocortisone and heparin. Some scaffolds had been reconstructed with macrophages, while others did not include them. On the day of infection, scaffolds were washed three times with pre-warmed (37°C) PBS to remove excess mucus. Infection was performed using IAV at a MOI of 1, based on an estimated 1.5×10^6^ epithelial cells per scaffold. Virus inoculum was applied on the apical side and scaffolds were incubated with the virus for 1h at 37°C in a humidified atmosphere with 5% CO_2_, after which they were returned to air-liquid interface conditions over 3mL of PneumaCult-ALI medium. Mock-infected controls were treated identically but incubated with PBS instead of viruses. Supernatants were collected at 6h and 48h post-infection to quantify viral titers by plaque assay, and at 6h, 24h, and 48h to assess cytokine release. Infected mucosa models were harvested at 6h and 48h post-infection for virus detection by immunofluorescence and RNA extraction for RT-qPCR analysis.

### 2.12 Immunofluorescence microscopy

Cells were seeded on poly-L-lysine-coated Lab-Tek chamber slides (Thermofisher) at a density of 2□×□10^5^ cells per well and incubated for 1h at 37°C in a humidified atmosphere with 5% CO_2_. Cells, mid-turbinate tissues, and reconstructed nasal models were fixed with 4% paraformaldehyde (PFA) (Sigma-Aldrich) for 30min at room temperature, followed by a wash with PBS. Tissues were incubated in 30% sucrose for 24h at 4°C before embedding in OCT (Cell Path). Sections were cut at 30μm thickness. Cells and tissue sections were permeabilized and blocked for 30min at room temperature in immunofluorescence staining buffer consisting of PBS supplemented with 5% BSA, 0.1% Triton X-100 (Sigma-Aldrich), and 2% normal horse serum (Jackson ImmunoResearch). Samples were incubated with primary antibodies diluted in the same buffer for 3h at room temperature, washed three times with PBS, and subsequently incubated with secondary antibodies and DAPI for 1h at room temperature. After three additional PBS washes, a 30min incubation with 2μg/mL streptavidin AF488 (Molecular Probes) diluted in blocking buffer was performed for samples stained with biotin-conjugated antibodies. Samples were washed again three times with PBS and mounted using Fluoromount-G™ mounting medium (Thermofisher). Imaging was performed using a spinning disk inverted microscope (Zeiss) equipped with a Yokogawa CSU confocal head and controlled by Metamorph software (Molecular Devices). Image analysis was conducted using FIJI software^18^.

The antibodies used to detect epithelial and basal cells in nasal brushings, mid-turbinate tissues, and reconstructed nasal mucosa were: goat anti-human cytokeratin-19 (CK19) (R&D Systems), rat anti-human p63 (Biolegend), donkey anti-goat AF647 (Jackson ImmunoResearch), and donkey anti-rat-biotin (Jackson ImmunoResearch).

To detect differentiated epithelial cells in mid-turbinate tissues and nasal mucosa models, the following antibodies were used: mouse anti-human acetylated tubulin (Sigma-Aldrich), rabbit anti-human MUC5B, donkey anti-mouse AF555 (Thermofisher), and donkey anti-rabbit AF488 (Thermofisher).

To identify fibroblasts in the lamina propria of mid-turbinate tissues and nasal mucosa models, mouse anti-human vimentin (Biolegend) and donkey anti-mouse AF555 were used. Macrophages in the lamina propria were detected using: mouse anti-human CD68-PE (Biolegend), and donkey anti-mouse AF555 (Thermofisher)

For detection of IAV in infected reconstructed nasal mucosa, rabbit anti-IAV nucleoprotein (NP) (Abcam) and donkey anti-rabbit AF488 (Thermofisher) were used.

### 2.13 Virus titration by plaque assay

MDCK cells were cultured in DMEM medium supplemented with 10% FBS, 100U/mL of penicillin/streptomycin, 50μg/mL of gentamicin, and maintained at 37°C in a humidified atmosphere with 5% CO□. For plaque assays, confluent MDCK monolayers in 6-well plates were infected with 180μL of 10-fold serial dilutions of virus containing supernatants in PBS with Ca^2^□/Mg^2^□ and 0.2% BSA. Virus adsorption was carried out for 1h at 37°C with gentle rotation every 10min. After removal of the virus suspension, cells were overlaid with 2mL of Minimum Essential Medium (MEM) (Gibco) containing 1.8% agarose (Sigma-Aldrich) and 1μg/mL TPCK-treated trypsin. Plates were incubated for 72h at 37°C. After incubation, cells were fixed with 4% PFA for 30min at room temperature and stained with 2% crystal violet (Sigma-Aldrich) in 20% ethanol. Plaques were manually counted, and virus titers were expressed as plaque-forming units per milliliter (PFU/mL).

### 2.14 Gene expression analysis by RT-qPCR

Total RNA was extracted from infected nasal mucosa models at 6h and 48h post-infection, using the NucleoZOL kit (Macherey-Nagel) according to the manufacturer’s instructions. Genomic DNA was removed with the RapidOut DNA Removal Kit (Thermo Fisher Scientific), and cDNA was synthesized using the RevertAid First Strand cDNA Synthesis Kit (Thermofisher). qPCR reactions were prepared in 10μL volumes using PowerUp SYBR Green Fast Master Mix (Thermofisher), 200nM of each primer (sequences listed in **Supplementary Table 2**) and 20ng of cDNA per reaction. Amplification was performed on a StepOnePlus™ Real-Time PCR System (Applied Biosystems) using the following cycling conditions: initial denaturation at 95°C for 2min, followed by 40 cycles of denaturation at 95°C for 15s and annealing/extension at 60°C for 30s. Relative gene expression, including viral and host interferon-related transcripts, was calculated using the 2^^−ΔΔCt^ method, where Ct values were first normalized to the endogenous control gene *ACTB* (ΔCt), and then to the mock-treated models without macrophages (ΔΔCt). Data was obtained from four independent experiments.

### 2.15 Quantification of cytokines in culture supernatants

Supernatants from infected mucosa models were collected at 0h, 6h, 24h, and 48h post-infection. Cytokine levels were measured using the LEGENDplex™ Hu Anti-Virus Response Panel 1 (13-plex) (Biolegend) according to the manufacturer’s instructions. Supernatants were diluted 1:1024 for detection of IL-6 and IL-8, and 1:4 for detection of IFN-λ1, IL-1β, IFN-λ2, granulocyte-macrophage colony-stimulating factor (GM-CSF), and IFN-β. Data acquisition was performed using a Beckman Coulter CytoFLEX LX flow cytometer and analysis was conducted with the Qognit LEGENDplex™ Data Analysis Software.

### 2.16 Statistical analysis

Statistical analyses were performed using GraphPad Prism version 8.0. Differences between groups were assessed using the Kruskal-Wallis test followed by Dunn’s multiple comparisons test. A *p*-value < 0.05 was considered statistically significant.

## 3. Results

### 3.1 Isolation and expansion of primary cells from the human nasal mucosa

In order to generate a reconstructed 3D tissue model of the human respiratory mucosa in the upper airways, different primary cell types were isolated from human donors: fibroblasts as the main element of the lamina propria, macrophages as innate immune sentinels, and epithelial basal cells able to differentiate into a respiratory epithelium. To obtain nasal fibroblasts, we digested biopsies from the mid-turbinate area of the nasal cavity. The single-cell suspension derived from the digested tissue was a mixture of fibroblasts (EpCAM− CD31− CD90+), epithelial cells (EpCAM+) endothelial cells (CD31+) and hematopoietic cells (CD45+). After 7 days in culture, over 90% of the expanded cells were identified as CD90+ nasal fibroblasts **(Figure 1A, B)**.

**Figure 1.**
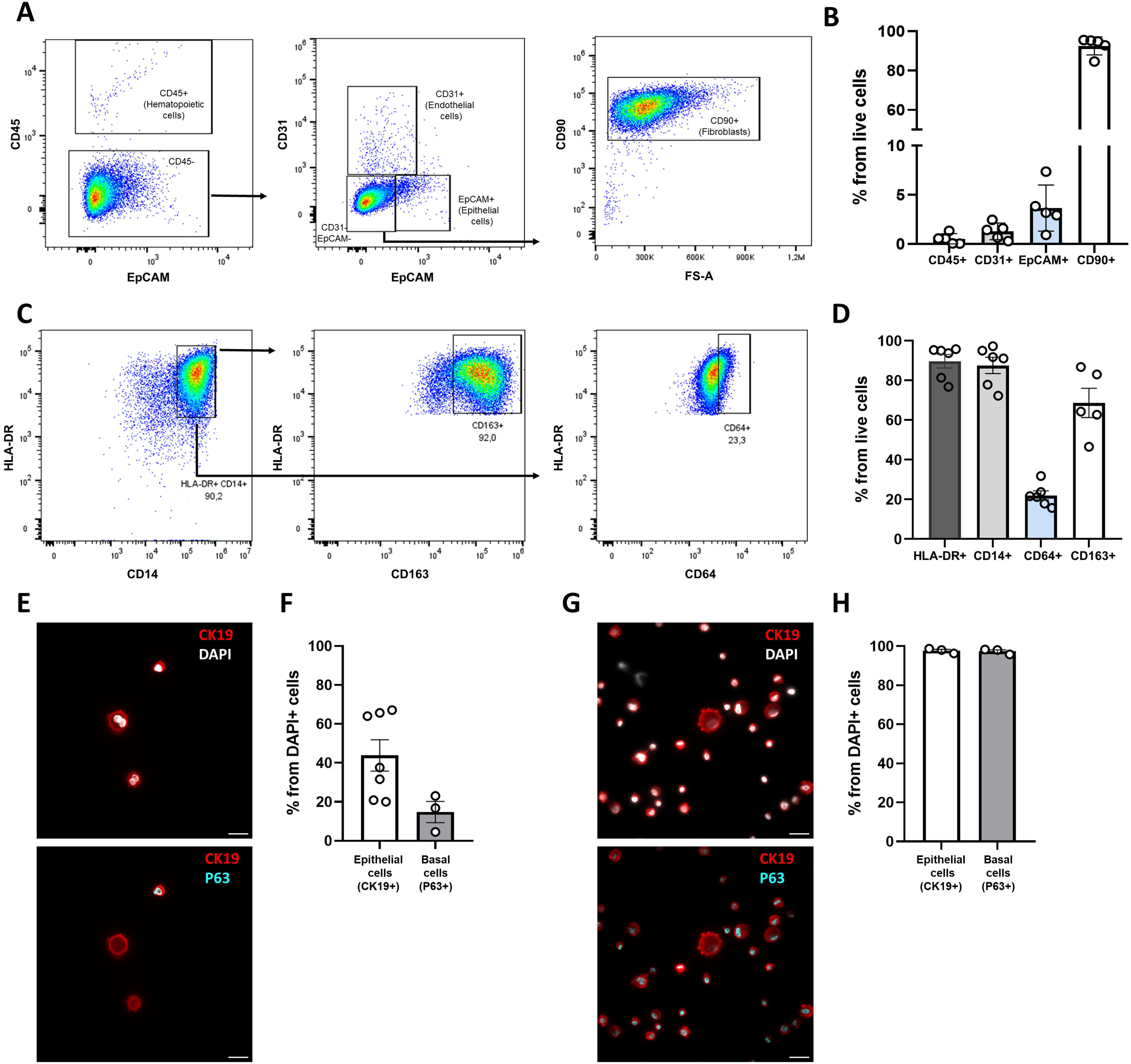
Amplification of human primary cells to construct an immunocompetent 3D model of the respiratory mucosa. **(A-B)** Characterization of stromal cells isolated from the nasal mucosa. **(A)** Representative flow cytometry plot of live single-cell suspensions from digested mid-turbinate biopsies, after 1 week of culture in complete DMEM, and **(B)** quantification of cell subsets (CD45+ hematopoietic cells, CD31+ endothelial cells, EpCAM+ epithelial cells and CD90+ fibroblasts). **(C-D)** Characterization of macrophages differentiated from blood monocytes, after 1 week in complete RPMI medium supplemented with M-CSF. **(C)** Representative flow cytometry plots depicting the expression of different macrophage markers (HLA-DR, CD14, CD163, CD64) and **(D)** quantification of subset proportions. **(E-H)** Characterization of epithelial cells. **(E, G)** Representative immunofluorescent staining of **(E)** freshly isolated cells from nasal brushings and **(G)** cells amplified in PneumaCult-Ex Plus medium after passage 3. Scale bars=20μm. **(F, H)** Proportions of CK19+ epithelial cells and p63+ basal cells among DAPI+ total cells in **(F)** freshly isolated and **(H)** cultured cells. Data in bar graphs are presented as mean ± SEM. Each data point represents one independent experiment. CK19, Cytokeratin-19; M-CSF, Macrophage Colony Stimulating Factor.

Macrophages were differentiated from blood monocytes using M-CSF. After 7 days, the cells were mostly adherent and consistently displayed macrophage markers **(Figure 1C, D)**. Over 80% of live cells co-expressed HLA-DR and CD14. In addition, around 60% of the differentiated cells were CD163+ and 20% CD64+ **(Figure 1D)**, with an overlap between the two populations, confirming their macrophage identity.

Most of the upper airway epithelium is composed of terminally differentiated epithelial cells, like ciliated cells and mucus-producing goblet cells. Basal cells (p63+) are located in the innermost layer of the epithelium and function as progenitors, capable of differentiating into different specialized epithelial cell populations of the upper airways. Brushings from the nasal cavity of human donors resulted in a cell suspension containing roughly between 20% to 70% of EpCAM+ epithelial cells **(Figure 1E, F),** while the proportion of basal cells recovered from the brushings was less than 20% of the total **(Figure 1E, F)**. However, after culturing the cell suspensions in PneumaCult-Ex Plus, a medium supporting the growth of bronchial epithelial cells, the cultures appeared almost entirely composed of basal cells after 3 passages **(Figure 1G, H)**.

### 3.2 The reconstructed respiratory mucosa model closely resembles the human nasal mucosa

To construct the 3D in vitro model, we used a mixture of type I bovine collagen and chitosan as a scaffold. When lyophilized, this turned into a porous structure which can be kept for long-term storage. Upon rehydration, the scaffold mimicked the role of the ECM, offering anchoring support for the cells included in the lamina propria of the 3D model. In the respiratory mucosa, fibroblasts are the most represented cells in the lamina propria, where they are the main producers of ECM components. Thus, we initially used nasal fibroblasts to colonize the collagen-chitosan matrices. They were seeded on both sides of the scaffolds and cultured for 6 weeks to allow full colonization and secretion of additional ECM for better support of the epithelium. Vitamin C was provided into the culture medium to further stimulate collagen production by fibroblasts. On week 6 of the scaffold colonization by fibroblasts, macrophages were introduced into the model. Finally, to obtain a fully differentiated epithelium, we seeded respiratory basal cells, expanded for <3 passages, on the top side of the scaffolds and allowed their expansion for 1 week. To induce their differentiation, the scaffolds were lifted at the air-liquid interface for 6 additional weeks. The experimental approach to build a human respiratory mucosa model is depicted in **Figure 2A**.

**Figure 2.**
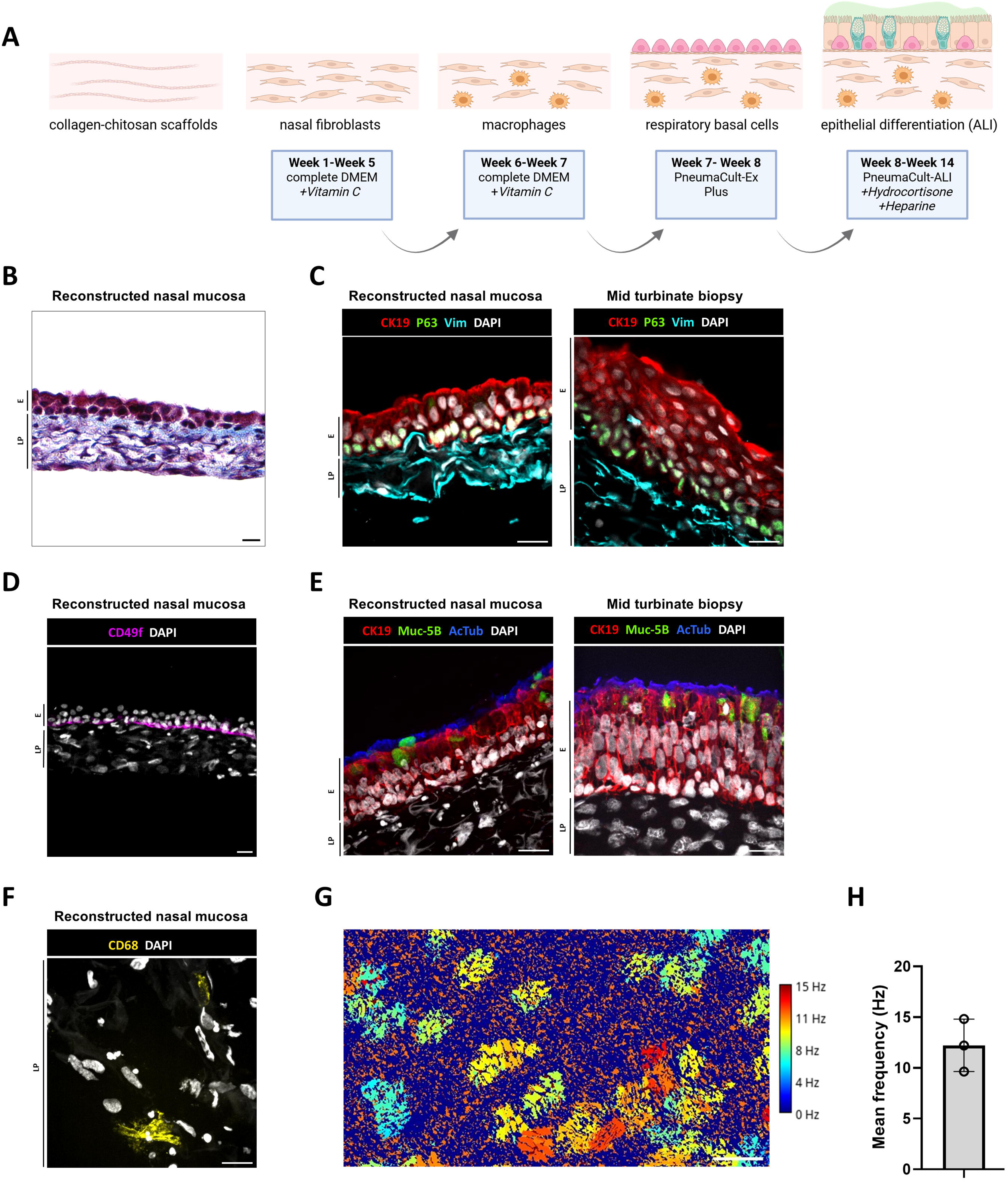
The immunocompetent 3D model replicates key features of the human nasal mucosa. **(A)** Experimental workflow for the construction of the model. Nasal fibroblasts were seeded in collagen-chitosan scaffolds and cultured in complete DMEM-F12 medium supplemented with vitamin C. Macrophages were introduced after 6 weeks. Basal cells expanded from nasal brushings were seeded on week 7 and the scaffold was cultured for 1 week in PneumaCult-Ex Plus medium. Sponges were lifted to the air-liquid interface on week 8 and cultured with PneumaCult-ALI medium supplemented with hydrocortisone and heparin until week 14. Illustration created with Biorender. **(B)** Presence of an epithelium (purple) and lamina propria with cellular components (purple) and collagen (blue). **(C, E)** Comparison of the respiratory mucosa model with mid-turbinate biopsies. **(C)** Presence of an organized epithelium (CK19+) containing basal cells (p63+) and a lamina propria formed by fibroblasts (Vim+). **(D)** Presence of a basal lamina revealed by the CD49f/α6-integrin staining of basal epithelium in the mucosa model. **(E)** Differentiation into a functional respiratory epithelium containing ciliated cells (AcTub+) and goblet cells (Muc-5B+). **(F)** Detection of macrophages (CD68+) in the lamina propria of the mucosa model. Section thickness = 30μm, scale bars = 20μm. Nuclei were stained with DAPI. CK19, Cytokeratin-19; Vim, vimentin; AcTub, acetylated tubulin; Muc-5B, mucin-5B; E, epithelium; LP, lamina propria. **(G, H)** Analysis of ciliary beating activity **(G)** Representative frequency map of ciliary beating. Fast Fourier transform analysis of time-lapse recordings was used to generate the map, with regions where pixels exhibit similar frequency values highlighted, corresponding to local cilia beating activity. **(H)** Average cilia beating frequency. Mean frequencies were measured for each image within segmented regions corresponding to cilia activity. Each data point represents the average for one mucosa model.

The reconstructed respiratory model showed an organized epithelium and a lamina propria with cellular components and collagen fibers **(Figure 2B)** We next examined how this architecture recapitulates features of the native tissue by comparing it with human mid-turbinate nasal mucosa biopsies. The 3D models showed a dense arrangement of fibroblasts expressing vimentin in the lamina propria, similarly to the mid-turbinate sections **(Figure 2C)**. The presence of an organized epithelium was also confirmed: it was composed of epithelial cells (CK19+) at the apical region, with progenitor basal cells, displaying nuclear p63 expression, in the innermost layer, right above the lamina propria **(Figure 2C)**. Basal cells also showed polarized expression of CD49f/α6-integrin at their basal domain, which permits their anchoring to the basement membrane **(Figure 2D)**. A functional respiratory epithelium is characterized by the presence of ciliated cells and mucus-producing goblet cells, that together form the muco-ciliary apparatus and expulse particles and pathogens trapped in mucus. The presence of ciliated cells was determined by staining for acetylated tubulin (AcTub) in the apical cilia, whereas goblet cells were identified by the expression of MUC5B, one of the most abundant mucins secreted in mucus **(Figure 2E)**. Macrophages were detected in the lamina propria by the expression of the pan-macrophage marker CD68 **(Figure 2F).**

Finally, cilia beating activity, which was visible at the surface of the differentiated epithelium, was quantified by time-lapse microscopy. Ciliary beating frequency maps were generated, highlighting regions where pixels exhibit similar frequency values **(Figure 2G)**. Mean frequencies were measured for each region with detectable activity. The average ciliary beating frequency was 12.2[Hz (**Figure 2H**), consistent with the reported range of 10-20 Hz for human airway cilia^20^.

Altogether, the respiratory mucosa model presented a fibroblast-rich lamina propria with integrated macrophages and a fully differentiated epithelium including basal, ciliated and goblet cells, organized in a similar way as the native human tissue.

### 3.3 Reconstructed respiratory mucosa models were susceptible and permissive for IAV infection

To assess whether 3D mucosa model could be used as a new tool to study human respiratory viral infections, these models, constructed with and without macrophages in the lamina propria, were topically exposed to IAV at week 7 post air-lifting. In the previous week, hydrocortisone and heparin, used to promote the differentiation of the airway epithelium, were removed from the culture medium to prevent any effect on immune responses or viral attachment, respectively. The presence of the virus in the reconstructed mucosa was assessed by immunofluorescence and RT-qPCR at early (6h) and late (48h) timepoints. At 48h post-infection, IAV nucleoprotein (NP) was detected in the differentiated ciliated epithelium by immunofluorescence **(Figure 3A)**. At 6h, the NP was not detectable at the protein level **(Figure 3A)**, yet NP transcription had already started to increase and reached significantly higher levels than in mock-infected models by 48h **(Figure 3B)**. To confirm that the replicative cycle was completed and infectious viral particles produced, virus titers in the culture supernatants were determined by plaque assay on MDCK cells. At 48h post-infection, viral titers reached between 10^5^ to 10^6^ PFU/mL, which was significantly higher than at the 6h early timepoint, indicating viral replication within the infected models **(Figure 3C)**. Notably, the presence of macrophages in the mucosa models did not affect viral replication **(Figure 3B, C)**.

**Figure 3.**
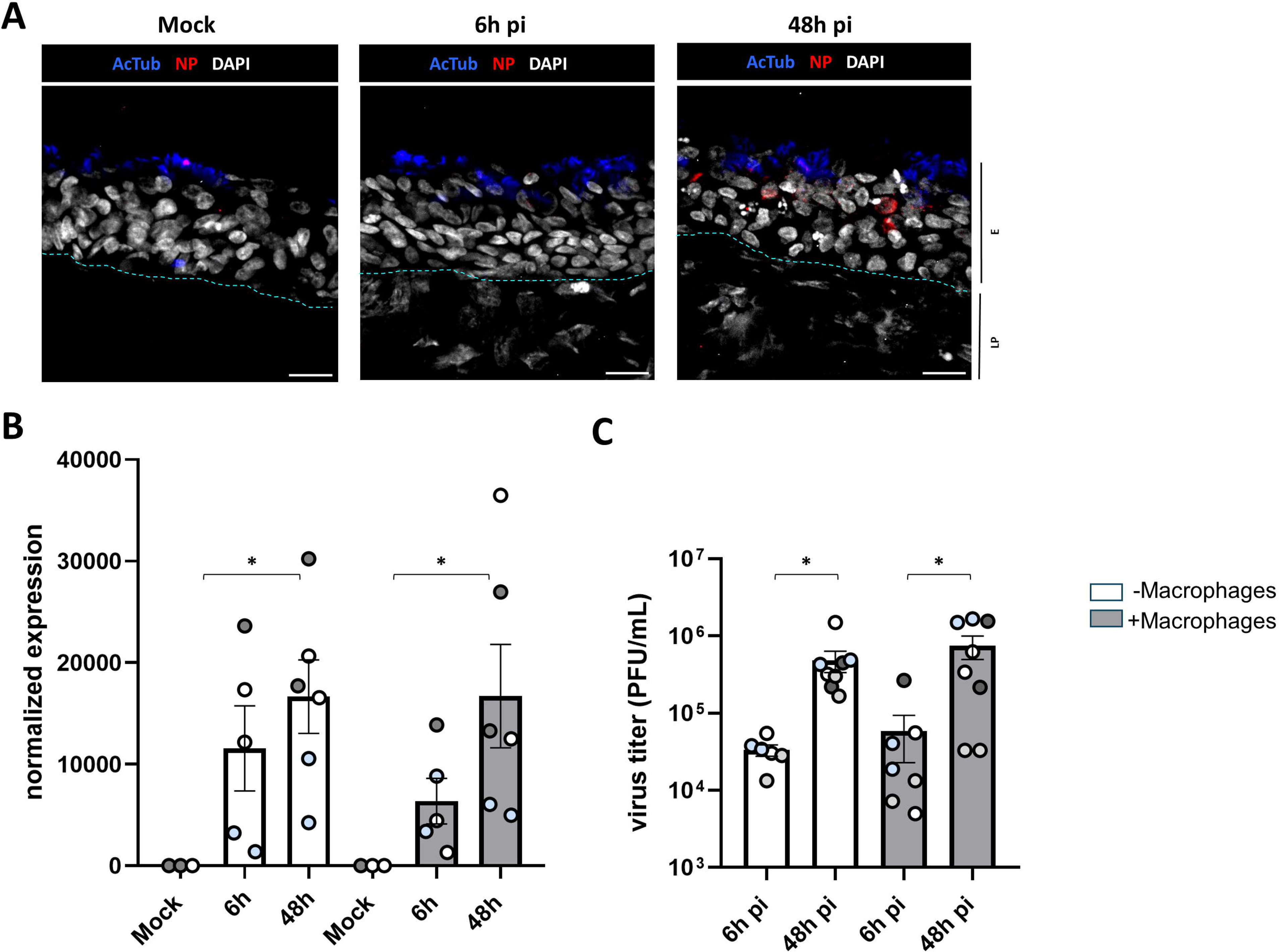
Influenza A virus productively infects human respiratory mucosa models. Human respiratory mucosa models, generated either with (+M) or without (-M) macrophages, were infected with influenza A virus (IAV) at a multiplicity of infection (MOI) of 1. Infection was assessed at early (6h) and late (48h) timepoints post-infection (pi), or in mock-infected controls. **(A)** Immunofluorescent staining of infected mucosa models. Viral presence is indicated by IAV nucleoprotein (NP). Differentiated respiratory epithelium is labeled with acetylated tubulin (AcTub) and cytokeratin-19 (CK19), and nuclei are stained with DAPI. Section thickness = 30□μm, scale bars = 20□μm. Dashed lines depict the limits of the epithelium (E) and the lamina propria (LP). **(B)** Expression of mRNA encoding IAV NP in infected mucosa models, detected by RT-qPCR. The relative expression levels were calculated using the 2^−ΔΔCt^ method, normalized to the internal control gene *ACTB*, with the non-infected group without macrophages (mock −M) serving as the reference sample. **(C)** Viral titers in the supernatants of infected respiratory mucosa models, measured by plaque assay on MDCK cells. Titers are expressed as plaque-forming units per milliliter (PFU/mL). Data represents results from 4 independent experiments. Bar graphs are presented as mean ± SEM, with each data point representing one mucosa model and colors denoting the different experiments. Statistical significance was determined using the Kruskal-Wallis test followed by Dunn’s multiple comparison test. Only significant differences are shown: *, p□<□0.05.

### 3.4 Innate immune responses are triggered in infected mucosa models

The first event of an antiviral immune response in the respiratory mucosa consists in the engagement of pattern recognition receptors in epithelial, stromal, and immune cells, which leads to the production of proinflammatory cytokines, interferons (IFNs) and interferon-stimulated genes (ISGs) to control virus spread. Therefore, we assessed the induction of these mediators in the reconstructed respiratory mucosa upon IAV infection.

At 48h, culture supernatants exhibited an enhanced production of proinflammatory cytokines (IL-6, IL-8, GM-CSF and IL-1β) **(Figure 4)**. Type I IFN-β, as well as type III IFN-λ1 and IFN-λ2 were also steadily increased over the course of infection **(Figure 4)**. In models containing macrophages, we observed a trend towards augmented secretion of several soluble mediators (IFN-β, IFN-λ2, IL-6 and IL-8) or decreased secretion of IL-1β. However, these differences did not reach statistical significance **(Figure 4).** IFNB mRNA levels were augmented at 48h, consistent with the IFN-β protein levels detected in the supernatant, while IFNA2 remained unchanged **(Figure 5)**. In line with this, mRNA expression of several ISGs (ISG15, RSAD2, MX1 and OAS1) was significantly increased at 48h post-infection, whereas DDX58 did not show any change **(Figure 5)**.

**Figure 4.**
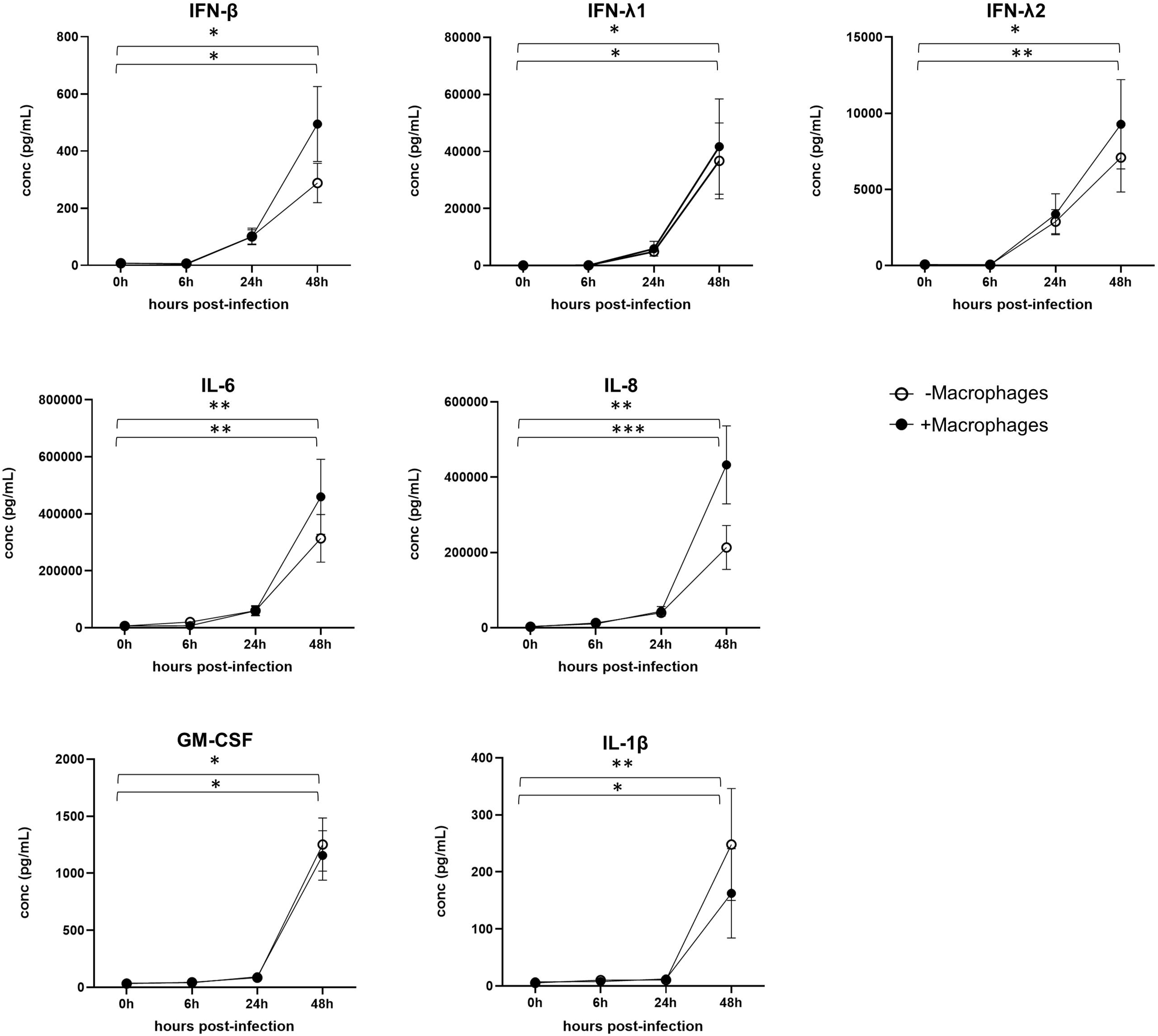
Influenza A infection promotes the secretion of proinflammatory and antiviral cytokines. Human respiratory mucosa models, generated either with (+M) or without (-M) macrophages, were infected with influenza A virus at a multiplicity of infection (MOI) of 1. Secretion of cytokines in the supernatant was evaluated at early at early (0h and 6□h) and late (24h and 48□h) timepoints post-infection (pi). Data represent results from 4 independent experiments. Bar graphs are presented as mean ± SEM, with each data point representing one mucosa model and colors denoting the different experiments. Statistical significance was determined using the Kruskal-Wallis test followed by Dunn’s multiple comparisons test. Only significant differences are shown: *, p□<□0.05; **, p□<□0.01; ***, p <□0.001. Upper brackets depict comparisons for mucosa models without macrophages, and lower brackets for models with macrophages. No significant differences were found between models with and without macrophages.

**Figure 5.**
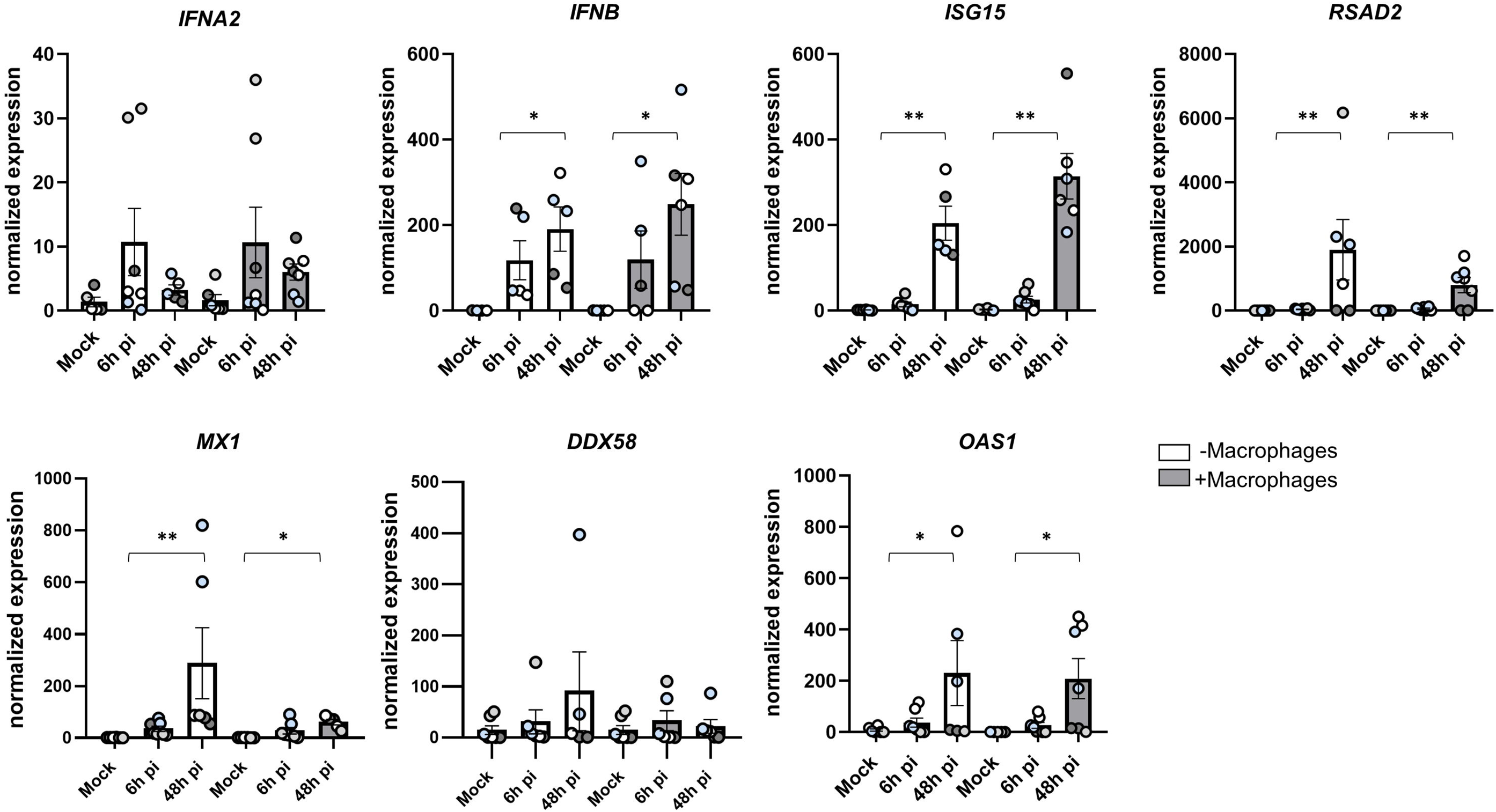
Influenza A infection triggers the expression of interferon-related genes. Human respiratory mucosa models, generated either with (+M) or without (-M) macrophages, were infected with influenza A virus at a multiplicity of infection (MOI) of 1. Expression levels of interferon-related genes were evaluated by RT-qPCR at early (6h) and late (48h) timepoints post-infection (pi), or mock-infected. The relative expression levels were calculated using the 2^−ΔΔCt^ method, normalized to the internal control gene *ACTB*, with the non-infected group without macrophages (mock −M) serving as the reference sample. Data represent results from 4 independent experiments. Bar graphs are presented as mean ± SEM, with each data point representing one mucosa model and colors denoting the different experiments. Statistical significance was determined using the Kruskal-Wallis test followed by Dunn’s multiple comparison test. Only significant differences are shown: *, p□<□0.05; **, p□<□0.01.

## 4. Discussion

Human respiratory tissue models have increasingly gained popularity as advanced tools to study host-pathogen interactions in a physiologically relevant environment^7^. Among them, ALI cultures remain the most established system to recapitulate airway epithelial differentiation and barrier function^8,9^. However, they fail to fully capture the complexity of the tissue. Here, we developed a 3D model of the human respiratory mucosa including not only a fully differentiated airway epithelium, but also a supporting lamina propria that included key stromal and immune components.

The lamina propria in the respiratory mucosa is rich in collagen and elastin fibers, intertwined with glycosaminoglycans^21,22^. These components contribute to airway elasticity, water balance, cell proliferation and migration^21^. To replicate these features in vitro, we used type I collagen and chitosan to form an artificial ECM. Chitosan has cationic charges and can form water-insoluble ionic complexes with collagen molecules^23^. This combination results in a 3D scaffold that confers mechanical strength to the model and structural support to stromal cells, while preserving flexibility and hydrophilicity^24^. Moreover, we used chitosan with a high degree of deacetylation, which prevents the degradation of the matrix in aqueous medium^25^, supporting the extensive culture time required to establish the model.

As the main cellular component of the lamina propria, we included CD90+ nasal fibroblasts derived from nasal mid-turbinates. As shown by vimentin staining, they fully colonized the collagen-chitosan scaffolds, forming a lamina propria closely resembling the native human nasal mucosa. Fibroblasts do not only provide structural support to the tissue but also influence the differentiation and repair of the epithelium^26–28^ and actively contribute to ECM production and remodeling^27^. Airway fibroblasts are also major producers of components of the basement membrane, which facilitates the anchoring of epithelial cells^29^. In the hereby presented mucosa model, basement membrane formation was evidenced by expression of α6-integrin by the most basal epithelial cell layer, which mediates adhesion to the basement membrane.

Airway basal cells were expanded from nasal brushing suspensions and used as a starting point to differentiate a respiratory epithelium. In the upper airways, basal cells are the main progenitors, giving rise to the two most abundant epithelial cells: ciliated cells and goblet cells^30^. In the respiratory mucosa model, we observed the presence of both differentiated cell types, in the context of a pseudo-stratified airway epithelium. Basal cells (p63+) were aligned immediately above the lamina propria, with mucus-producing goblet cells (MUC5B+) and ciliated cells (AcTub+) located in the apical side. Moreover, ciliated cells exhibited active and coordinated cilia beating, consistent with physiological behavior. The nasal mucosa is the first point of entry for respiratory viruses. At this level, mucociliary clearance and innate immune responses act as immediate defense mechanisms to limit viral spread to the lower airways^31,32^, where most pathological reactions typically occur^33^. A significant part of the existing in vitro airway models aims at reproducing areas in the lower respiratory tract, notably bronchi or lungs. However, upper and lower airway epithelia differ in their cellular composition^34,35^, gene expression^35–37^, susceptibility to infection by certain respiratory viruses^38–40^, and the magnitude of their IFN-mediated responses^41^. By using primary cells derived from the nasal cavity, the resulting 3D model is expected to better recapitulate early interactions occurring at the initial site of infection.

Viral infections of the respiratory mucosa elicit robust type I and type III IFN responses, along with the secretion of pro-inflammatory cytokines and chemokines that orchestrate systemic effects and regulate the recruitment and activity of innate and adaptive immune cells^32^. To evaluate the capacity of the reconstructed nasal mucosa to mount an antiviral innate immune response, the models were infected with IAV. The models were permissive to the infection, and the increase of inflammatory cytokines, type I and type III IFN was consistent with an antiviral innate immune response. This was further supported by the upregulation of multiple ISGs known to restrict IAV infection^42,43^.

Macrophages are central components of innate immune responses, contributing to tissue repair, pathogen sensing and regulation of inflammatory responses in the context of respiratory infections^44^, including IAV^46^. In the steady state, resident macrophages derived from fetal precursors are predominant in the airways but upon viral infections, monocytes are recruited from the blood and derived into macrophages to further support local immune responses^45^. We introduced monocyte-derived macrophages produced in vitro into the lamina propria of the respiratory mucosa model to better mimic innate immune responses within the tissue. Their presence at the end of the culture was confirmed and they were found distributed across different depths of the model. Surprisingly, the addition of macrophages did not seem to significantly modulate the magnitude of the antiviral immune responses. This was in line with an unaltered detection of viral genome and particles and suggests that, in this experimental system, respiratory epithelial cells and fibroblasts were the principal sources of IFNs and cytokines. Further characterization of the viability and phenotype of macrophages at the time of infection, as well as optimization of culture conditions may be necessary to unlock their functional contribution and better mimic in vivo immune responses. Of note, we detected the production of GM-CSF, a factor which may influence the function of macrophages introduced into the model. Finally, tissue-resident macrophages may be better mimicked by induced pluripotent stem cell-derived macrophages, which display features of embryonically seeded macrophages in other organs^47^.

Altogether, we have developed an immunocompetent nasal mucosa model comprising a fully differentiated respiratory epithelium and a lamina propria that closely recapitulates the structural organization of the human upper respiratory tract and demonstrated its suitability for studies on early host-pathogen interactions in the context of viral infections. This model could represent a valuable platform for advancing our understanding of respiratory mucosal immunity and for guiding the development of novel preventive and therapeutic strategies against known and emerging respiratory viruses.

## Author contributions

A. Pérez-Riverón and V. Flacher conceptualized the research. C. Debry and L. Fath provided biopsies from consenting patients. A. Pérez-Riverón, V. Flacher, B. Voisin, F. Foisset, C. Lehalle, N. Frossard and J. de Vos contributed to establishing the 3D culture protocol. J. Li and R. Smyth provided viral strains and related reagents. A. Pérez-Riverón, E. Deiss, A. Alléon, P. Ateni and JD. Fauny performed experiments and analyzed the data. A. Pérez-Riverón and V. Flacher wrote the manuscript, with critical revisions by B. Voisin and CG. Mueller.

---

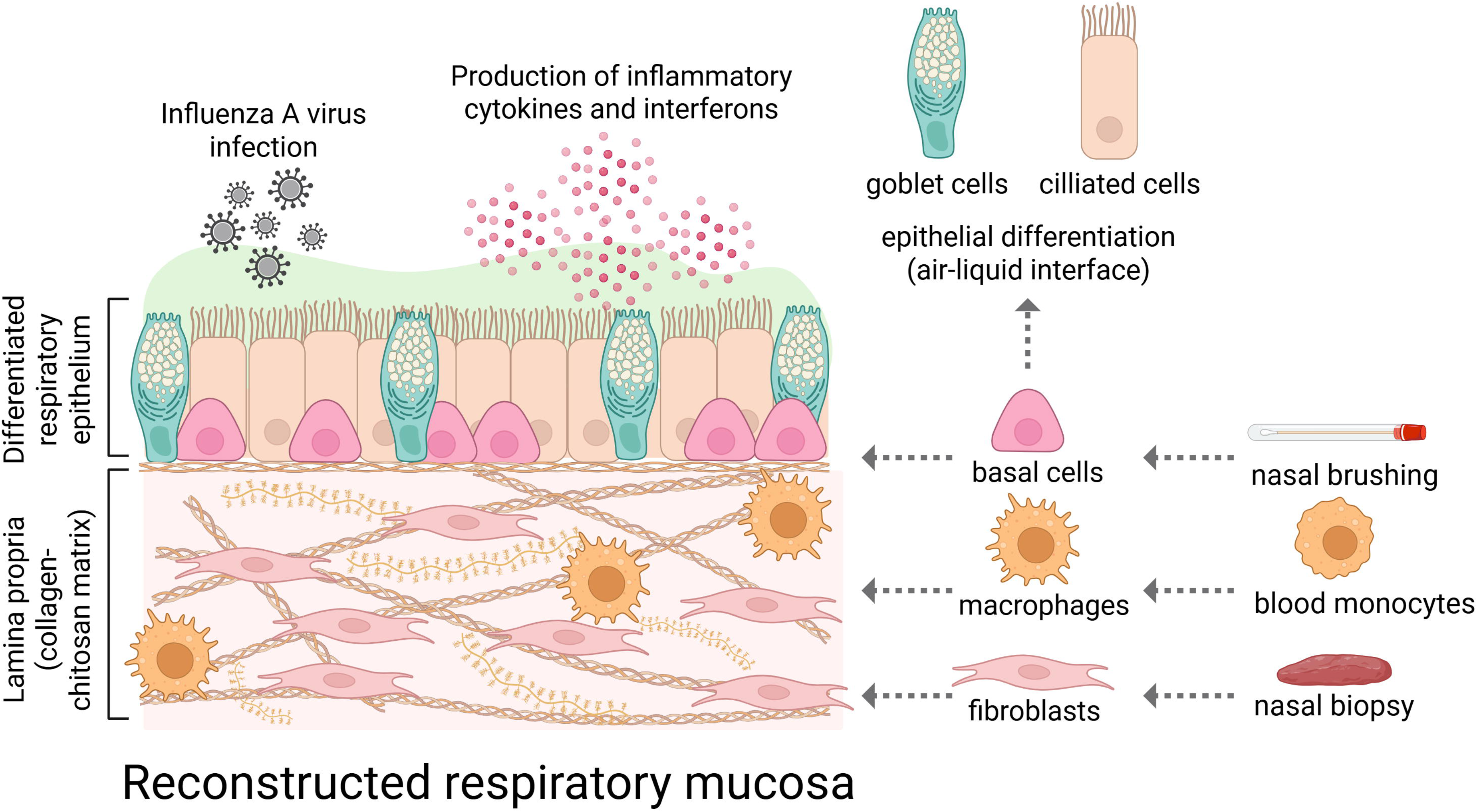

