## Supporting Information for "Influenza Virus Infection of an Immunocompetent Organotypic Model of the Human Respiratory Mucosa"

J. Li, R. Smyth

CNRS ARN/UPR9002 Architecture et Réactivité de l'ARN, Institut de Biologie Moléculaire et Cellulaire, Strasbourg, France

F. Foisset, C. Lehalle, N. Frossard

Laboratoire d’Innovation Thérapeutique, CNRS UMR7200, Laboratoire d’innovation Thérapeutique, Faculté de Pharmacie, University of Strasbourg, Illkirch, France

F. Foisset, J. de Vos

Institute for Regenerative Medicine and Biotherapy, INSERM U1183, University of Montpellier, Montpellier, France

J. de Vos

Department of Cell and Tissue Engineering, University of Montpellier, Montpellier, France

Hôpital Saint-Eloi, Montpellier, France

C. Debry, L. Fath

Ear-Nose and Throat and Head and Neck Surgery department, University Hospital of Strasbourg, Strasbourg, France

B. Voisin

University of Strasbourg, Strasbourg, France.

**Supporting information**

**Supplementary Table 1:** List of antibodies for flow cytometry and immunofluorescence stainings

| Target | Conjugate | Clone | Species | Concentration | Manufacturer | Reference |
| --- | --- | --- | --- | --- | --- | --- |
| EpCAM | AF488 | 158206 | recombinant human | 5μL/test | R&D | FAB9601G |
| CD31 | APC-Cy7 | WM59 | mouse | 2μg/mL | Biolegend | 303120 |
| CD90 | PE | 5E10 | mouse | 0.5μg/mL | Biolegend | 328110 |
| CD3 | APC | UCHT1 | mouse | 20μL/test | BD Biosciences | 555335 |
| CD45 | PE-Cy7 | HI30 | mouse | 5μL/test | Biolegend | 304016 |
| CD14 | PE | MEM-18 | mouse | 5μL/test | ImmunoTools | 21270144 |
| HLA-DR | AF700 | L203 | mouse | 10μL/test | R&D Systems | FAB4869 |
| CD64 | APC | 10.1 | mouse | 5μL/test | BD Pharmigen | 561189 |
| CD163 | FITC | 215927 | mouse | 5μL/test | R&D Systems | FAB1607G |
| CK19 | - | polyclonal | goat | 0.5μg/mL | R&D Systems | AF3506 |
| p63 | - | W17048F | rat | 2.5μg/mL | Biolegend | 699502 |
| MUC-5B | - | polyclonal | rabbit | 0.5μg/mL | Sigma-Aldrich | HPA008246 |
| AcTub | - | 6-11B-1 | mouse | 2.5μg/mL | Sigma-Aldrich | T7451 |
| vimentin | - | O91D3 | mouse | 2.5μg/mL | Biolegend | 677801 |
| CD68 | PE | Y1/82A | mouse | 0.12μg/mL | Biolegend | 333807 |
| IAV NP | - | polyclonal | rabbit | 2.5μg/mL | Abcam | Ab91648 |
| goat IgG | AF647 | polyclonal | donkey | 3.75μg/mL | Jackson ImmunoResearch | 705-605-147 |
| mouse IgG | AF555 | polyclonal | donkey | 5μg/mL | Thermofisher | A31570 |
| rabbit IgG | AF488 | polyclonal | donkey | 5μg/mL | Thermofisher | A21206 |
| rat IgG | biotin | polyclonal | donkey | 3.75μg/mL | Jackson ImmunoResearch | 712-065-153 |

**Supplementary Table 2:** List of primers for quantitative RT-PCR

| Name | Sequence (5’- 3’) |
| --- | --- |
| ACTB F | GCTCACCATGGATGATGATATCGC |
| ACTB R | CACATAGGAATCCTTCTGACCCAT |
| Influenza A NP F | AGTAGAAACAAGGGTATTTTTCCTTAATTGTCGTACTC |
| Influenza A NP R | GATGGAAGGTGCAAAACCAGAAGAAGTG |
| IFNA2 F | TGGGCTGTGATCTGCCTCAAAC |
| IFNA2 R | CAGCCTTTTGGAACTGGTTGCC |
| IFNB F | CTTGGATTCCTACAAAGAAGCAGC |
| IFNB R | TCCTCCTTCTGGAACTGCTGCA |
| ISG15 F | GAGAGGCAGCGAACTCATCT |
| ISG15 R | CTTCAGCTCTGACACCGACA |
| RSAD2 F | CCAGTGCAACTACAAATGCGGC |
| RSAD2 R | CGGTCTTGAAGAAATGGCTCTCC |
| MX1 F | GGCTGTTTACCAGACTCCGACA |
| MX1 R | CACAAAGCCTGGCAGCTCTCTA |
| DDX58 F | CACCTCAGTTGCTGATGAAGGC |
| DDX58 R | GTCAGAAGGAAGCACTTGCTACC |
| OAS1 F | AGGAAAGGTGCTTCCGAGGTAG |
| OAS1 R | GGACTGAGGAAGACAACCAGGT |
